# *GIGANTEA* accelerates wheat heading time through gene interactions converging on *FLOWERING LOCUS T1*

**DOI:** 10.1101/2023.07.11.548614

**Authors:** Chengxia Li, Huiqiong Lin, Juan M. Debernardi, Chaozhong Zhang, Jorge Dubcovsky

## Abstract

Precise regulation of flowering time is critical for cereal crops to synchronize reproductive development with optimum environmental conditions, thereby maximizing grain yield. The plant specific gene *GIGANTEA* (*GI*) plays an important role in the control of flowering time, with additional functions on the circadian clock and plant stress responses. In this study, we show that *GI* loss-of-function mutants in a photoperiod sensitive tetraploid wheat background exhibit significant delays in heading time under both long-day (LD) and short-day (SD) photoperiods, with stronger effects under LD. However, this interaction between GI and photoperiod is no longer observed in isogenic lines carrying either a photoperiod insensitive allele in the *PHOTOPERIOD1* (*PPD1*) gene or a loss-of-function allele in *EARLY FLOWERING 3* (*ELF3*), a known repressor of *PPD1.* These results suggest that the normal circadian regulation of *PPD1* is required for the differential effect of *GI* on heading time in different photoperiods. Using crosses between mutants or transgenic of *GI* and those of critical genes in the flowering regulation pathway, we show that *GI* accelerates wheat heading time by promoting *FLOWERING LOCUS T1* (*FT1*) expression via interactions with *ELF3, VERNALIZATION 2* (*VRN2*), *CONSTANS* (*CO*), and the age-dependent microRNA172-*APETALA2* (*AP2*) pathway, at both transcriptional and protein levels. Our study reveals conserved *GI* mechanisms between wheat and Arabidopsis, but also identifies specific interactions of GI with the distinctive photoperiod and vernalization pathways of the temperate grasses. These results provide valuable knowledge for modulating wheat heading time and engineering new varieties better adapted to a changing environment.

## Introduction

*GIGANTEA* (*GI*) is a plant-specific gene that encodes a large protein (1,173 amino acids in Arabidopsis) with no domains of known biochemical function (Fowler et al. 1999). The GI protein is predominantly localized to the nuclei where it forms nuclear bodies (Huq et al. 2000). Both *GI* transcription and GI protein abundance are regulated by the circadian clock (Fowler et al. 1999; Park et al. 1999; David et al. 2006) and can be detected throughout plant development (Fowler et al. 1999). In Arabidopsis plants grown under long-day photoperiod (16 h light/8 h dark, henceforth LD), the *GI* transcripts peak at Zeitgeber time 10 (ZT10); while under short-day (8 h light/16 h dark, henceforth SD), the peak is at ZT8 (Fowler et al. 1999; Park et al. 1999).

Defects in several components of the circadian clock have been shown to affect the *GI* transcription profile (Fowler et al. 1999; Lu et al. 2012). Chromatin Immunoprecipitation (ChIP) analysis showed that the evening complex (EC), including EARLY FLOWERING 3 (ELF3), EARLY FLOWERING 4 (ELF4) and LUX ARRYTHMO (LUX), regulates *GI* transcription by direct binding to its promoter (Ezer et al. 2017). ELF3 can also regulate GI protein degradation through its interaction with CONSTITUTIVELY PHOTOMORPHOGENIC1 (COP1) in an EC-independent manner (Yu et al. 2008). The Arabidopsis *elf3 gi* double mutant is unable to synchronize the circadian oscillator to the light-dark cycles, indicating that these two genes are essential for light entrainment of the clock (Anwer et al. 2020).

The multiple *gi* mutants identified in Arabidopsis (Rédei, 1962; Koornneef et al. 1991) show different effects on the length of the circadian clock period, but all of them exhibit a late flowering phenotype (Fowler et al. 1999; Park et al. 1999). The effect of *GI* on Arabidopsis flowering time is stronger under LD than under SD, indicating an interaction between *GI* and the photoperiod pathway (Araki and Komeda, 1993; Mizoguchi et al. 2005). In addition to its role in the circadian clock regulation and photoperiodic flowering response, *GI* has been shown to play important roles in multiple other physiological processes such as light signaling, chlorophyll accumulation, starch accumulation and stress tolerance (reviewed in Mishra & Panigrahi, 2015; Brandoli et al. 2020). Because GI can physically interact with many proteins of the circadian clock and photoperiodic flowering pathways, GI has been suggested to function as a scaffold or hub protein and orchestrates other protein interactions (Imaizumi and Kay, 2006).

In Arabidopsis, GI regulates the expression of *FT* and flowering time through both CONSTANS (CO)-dependent and CO-independent mechanisms. In the CO*-*dependent pathway, GI acts between the circadian oscillator and the main photoperiod gene *CO* to accelerate flowering by promoting both *CO* and *FT* mRNA abundance (Mizoguchi et al. 2005). GI interacts with the FLAVIN-BINDING, KELCH REPEAR, F-BOX1 (FKF1) protein to form a complex that regulates the protein abundance of CYCLING DOF FACTOR 1 (CDF1), a transcriptional repressor of *CO*. The degradation of CDF1 protein elevates the transcription of *CO*, thereby promoting *FT* expression and flowering (Sawa et al. 2007). In addition, GI can interact with CO directly and indirectly through interactions with FKF1 and ZEITLUPE (ZTL) to regulate CO protein stability (Song et al. 2014; Hwang et al. 2019). Finally, the interaction between GI and SPINDLY (SPY), which is a negative regulator of gibberellin signaling, can inhibit SPY activity and promote *FT* transcription in a CO-dependent manner (Tseng et al. 2004).

GI also regulates *FT* expression in Arabidopsis through two CO-independent pathways. The first one involves microRNA172 (miR172), which inhibits the expression of flowering repressors *TARGET OF EAT 1* (*TOE1*) and *APETALA 2* (*AP2*) (Jung et al. 2007). In the second CO-independent pathway, GI activates *FT* transcription through direct binding to the *FT* promoter alone or in a protein complex with *FT* repressors such as SHORT VEGETATIVE PHASE (SVP), TEMPRANILLO 1 (TEM1) and TEM2 (Sawa and Kay, 2011).

In Arabidopsis, *CO* plays a central role in the photoperiodic response by inducing flowering under LD conditions (Valverde et al. 2004), and *CO* homologs from rice and sorghum have also been shown to play major roles in the photoperiodic control of flowering (Yano et al. 2000; Yang et al. 2014). However, in the temperate grasses, the duplicated *CO1* and *CO2* genes show limited effects on the photoperiodic regulation of flowering time in the presence of *PHOTOPERIOD1* (*PPD1*). In the absence of functional *PPD1* alleles in wheat (Shaw et al. 2020), or in the presence of hypomorphic *PPD1* alleles in barley (Alqudah et al. 2014), the effect of *CO1* on the acceleration of heading time under LD is stronger than in the presence of *PPD1*. In the temperate grasses, *PPD1* is the dominant gene in the photoperiodic pathway, showing strong differences in heading time between LD and SD even in the absence of functional copies of both *CO1* and *CO2* (Shaw et al. 2020). Although the interactions between *GI* and *CO* have been extensively studied in Arabidopsis (Mizoguchi et al. 2005; Sawa et al. 2007) and rice (Hayama et al. 2003), the interactions between *GI* and *PPD1* remain largely unexplored.

Another understudied aspect of the role of *GI* in the temperate cereals is its interaction with *VRN2*, a LD repressor of *FT1* with no known homologs in eudicots (Yan et al. 2004). *VRN2* encodes a protein comprising a zinc finger motif at the N-terminus and a CCT domain at the C-terminus (Yan et al. 2004), which is orthologous to the rice protein GRAIN NUMBER, PLANT HEIGHT, AND HEADING DATE 7 (GHD7) (Xue et al. 2008; Woods et al. 2016). In rice, the GI protein has been shown to interact *in vivo* with GHD7 and to regulate its stability (Zheng et al. 2019). Loss-of-function mutations in *GHD7* result in earlier flowering under LD in rice (Xue et al. 2008), and mutations in *VRN2* result in accelerated heading under LD and a spring growth habit in wheat (Yan et al. 2004; Distelfeld et al. 2009).

In this study, we show that *GI* loss-of-function mutants in wheat exhibit significant delays in heading time under both LD and SD. This delay is larger under LD than under SD in photoperiod sensitive wheats (PS), but the differences disappear in photoperiod insensitive (PI) wheats and *elf3* mutants. Moreover, we show that GI regulates *FT1* expression and heading time through its interactions with *ELF3*, *VRN2*, *CO*, and the age-dependent miR172-AP2 pathway, revealing both conserved and specific mechanisms for the temperate grasses.

## Results

### *GI* loss-of-function mutants delay heading time in wheat under LD photoperiod

From a sequenced EMS mutant population in the tetraploid spring wheat variety Kronos (Krasileva et al. 2017), we identified two truncation mutants for the *GI* A-genome homoeolog (henceforth, *A-401* and *A-2019*) and two truncation mutants for the B-genome homeolog (henceforth *B-2205* and *B-3825*, Fig. 1A). Line *A-401* has a splice site mutation that leads to the retention of intron 4 and generates a premature stop codon that eliminates 1,034 amino acids (89.5%) of the GI protein (total 1,155 amino acids). The other three mutant lines all have premature stop codon mutations: *A-2019* (Q801*), *B-3825* (Q41*) and *B-2205* (W754*).

**Fig. 1.**
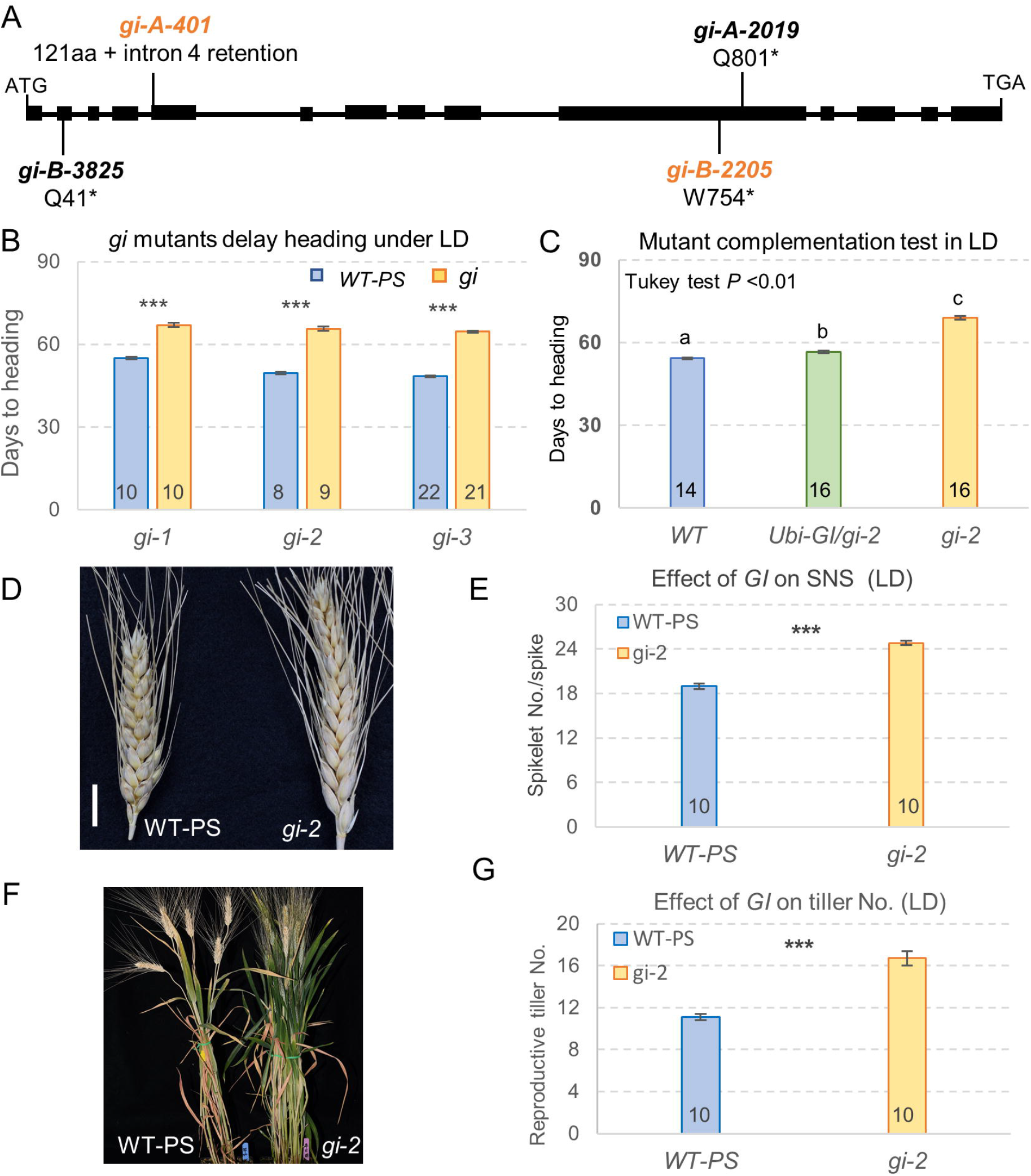
Effect of *gi* loss-of-function mutants and transgenic lines expressing *GI* under the maize ubiquitin promoter (*UBI-GI-MYC*) on heading time under long-day (LD). A, Schematic diagram showing *GI* gene structure and the positions of selected mutations. Exons are represented by rectangles and introns by a line. Mutant line *A-2019* was combined with *B-2205* to generate *gi-1*, *A-401* combined with *B-2205* to generate *gi-2*, and *A-401* combined with *B-3825* to generate *gi-3* (letters indicate the genomes, and numbers indicate the Kronos mutant identification). B, *gi* mutants in Kronos-PS show a significant delay in heading time relative to their control sister lines under LD. C, The *UBI-GI-MYC* transgene largely rescues the delayed heading time of *gi-2* mutant in Kronos-PS background under LD. Different letters above the bars indicate significant differences in Tukey test = *P* < 0.01. D-E, Kronos-PS spikes of *gi-2* mutants produce more spikelets per spike (SNS) than those of the wild type. The scale bar in D = 1 cm. F, Kronos-PS 12-week-old *gi-2* mutant compared to wild-type sister lines. G, The *gi-2* mutant plants produce more spike-bearing tillers than the control sister plants. Numbers inside bars indicate the number of plants included in each analysis. Error bars are standard errors of the means. *** = *P* < 0.001 for differences with the wild-type control using two-tailed *t-*test in B, E and G. Raw data and statistical analyses are available in Supplemental Data S1.

Kronos carries the photoperiod insensitive allele *Ppd-A1a*, which has a 1,027 bp deletion in the promoter region and is associated with misexpression of *PPD1*, accelerated induction of *FT1* expression, and early flowering under SD (Wilhelm et al. 2009). To study the effect of the *GI* mutations in a photoperiod sensitive background, we crossed the four *gi* mutants twice with a near-isogenic Kronos line carrying the photoperiod sensitive *Ppd-A1b* allele (henceforth Kronos-PS, Pearce et al. 2017), and intercrossed them to generate three different BC_1_F_2_ lines homozygous for *gi* mutants in both the A-and B-genome homoeologs. The *gi-1* line, generated from the cross between *A-2019* and *B-2205*, headed 12 d later than Kronos-PS, whereas the *gi-2* (*A-401* x *B-2205*) and *gi-3* (*A-401* x *B-3825*) mutants headed 16 d later (Fig. 1B). These results indicate that *GI* functions as a promoter of heading time in wheat under LD.

To further explore the function of *GI*, we generated wheat transgenic lines constitutively expressing a fusion of the full-length coding region of the *GI* gene and a C-terminal 4xMYC tag under the control of the maize *UBIQUITIN* promoter in a Kronos-PI background (hereafter referred to as *UBI-GI-MYC*). Among the six independent transgenic events, transgenic line #13 showed the highest transcript levels of *GI* relative to the non-transgenic sister line (Fig. S1A, *P* < 0.001) and was selected for further characterization under both LD and SD conditions.

Transgenic T_1_ plants of the *UBI-GI-MYC* line #13 headed 3.3 d earlier than the non-transgenic Kronos-PI controls when grown under LD (Fig. S1B), but showed no difference in heading time relative to the controls under SD (Fig. S1C). These results confirmed that *GI* is a LD promoter of wheat heading time.

We next crossed the *UBI-GI-MYC* #13 transgenic plant (PI) with the *gi-2* mutant (PS) to test the ability of the transgene to rescue the delayed heading time of the *gi-2* mutant. In the F_2_ progeny, we selected sister lines homozygous for the wild-type control (WT), the non-transgenic *gi-2* mutant (*gi-2*), and the combined *UBI-GI-MYC* / *gi-2* (all homozygous for the *PPD1-A1b* allele, PS). The *UBI-GI-MYC* / *gi-2* plants headed 12.4 d earlier than the *gi-2* mutant, but still 2.4 d later than the wild type under LD (Fig. 1C). These results indicate that the *UBI-GI-MYC* #13 transgene can largely, but not completely, rescue the delayed heading time of the *gi-2* mutant (Fig. 1C).

In addition to the late heading under LD, the *gi-2* mutants also showed an increased number of spikelets per spike (Fig.1D-E) and tillers (Fig. 1F-G). Compared with the wild-type control plants, the *gi-2* mutant produced on average 5.8 more spikelets per spike (Fig.1E) and had an average of 5.6 more spike-bearing tillers (Fig. 1G), likely associated with its extended vegetative phase.

### *GI* shows strong interaction with photoperiod in Kronos-PS, but not in Kronos-PI

When grown under SD (8 h light), Kronos-PI plants head between 80 and 100 days, whereas Kronos-PS plants fail to head (Shaw et al. 2020; Alvarez et al. 2023). Under these conditions Kronos-PI plants produce normal spikes and viable grains, whereas Kronos-PS plants undergo spike and stem elongation arrest, and spikes fail to emergence from the leaf sheaths (Pearce et al. 2017; Shaw et al. 2020). Similarly, the *gi-2* mutants failed to head under SD before the experiment was terminated at 150 d (Fig. 2A), but showed fully developed young spikes wrapped inside the leaf sheath (Fig. 2B).

**Fig. 2.**
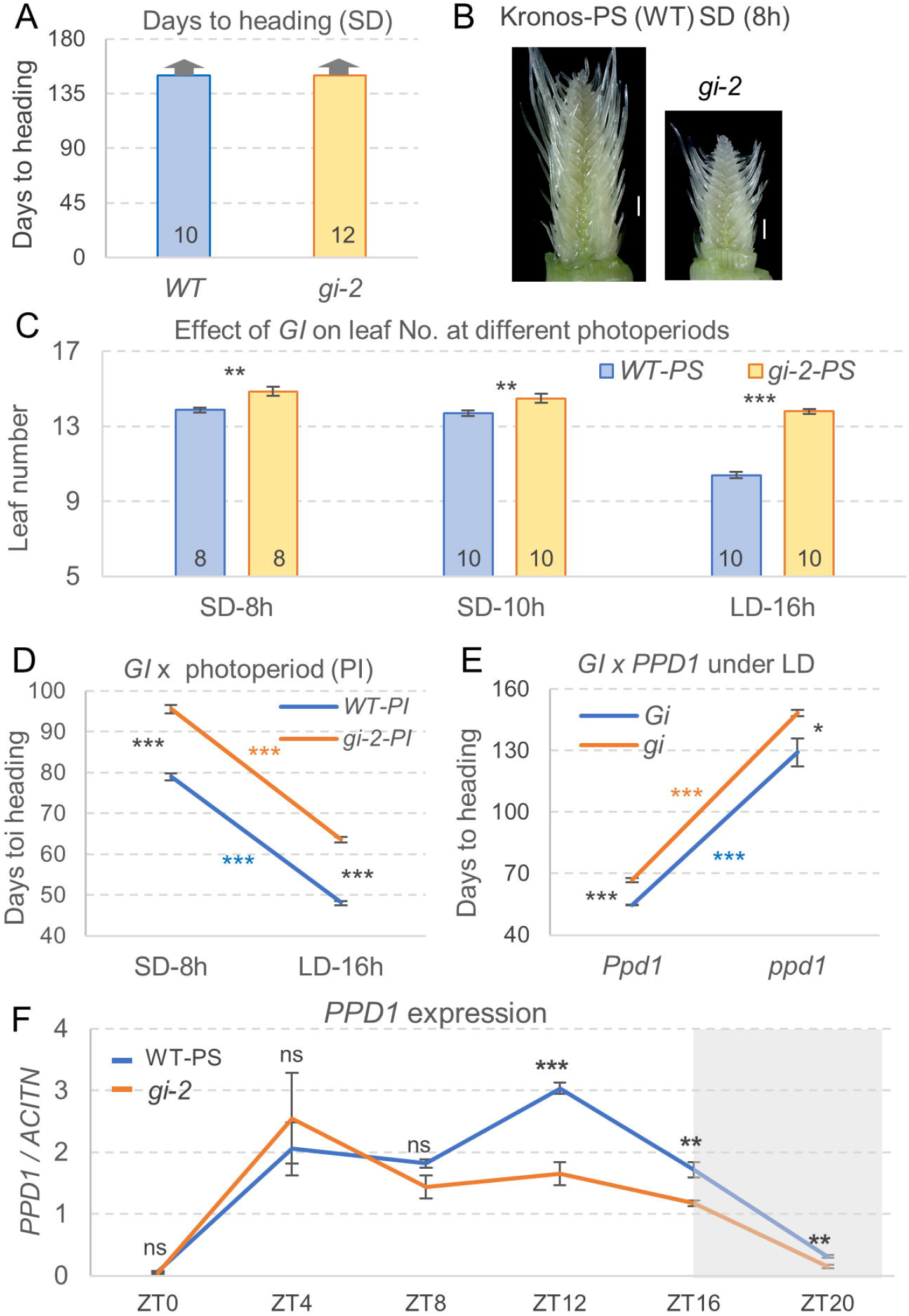
Interactions between *GI* and photoperiod in Kronos-PS, Kronos-PI and *ppd1* mutants. A, Heading time evaluation of Kronos-PS and *gi-2* mutant grown under SD (8h light). Arrows on top of bars indicate that both genotypes failed to head before the experiment was terminated at 150 d. B, Pictures of developing young spikes of Kronos-PS and *gi-2* mutant dissected 100 d after germination from the same batch of plants grown under SD as in A. Scale bar = 1000 µm. C, Effect of *gi-2* on leaf number under SD-8h, SD-10h and LD-16h in a Kronos-PS background. Numbers inside the bars in A and C indicate the number of plants included in the analysis for each genotype. D, Interaction graph showing the effect of *gi-2* on days to heading under SD-8h and LD-16h in a Kronos-PI background (n = 10 - 11). E, Interaction graph showing the effect of *gi-2* and *ppd1* mutants on heading time under LD. Error bars are standard errors of the means based on n = 7 - 8. F, Effect of *gi* mutations on *PPD1* transcription profiles under LD. The gray shaded area indicates the 8 h of darkness. Primers used in the qRT-PCR amplify both *PPD1* homeologs. The 4^th^ fully expanded leaves were collected at 4 h intervals for 24h. Expression levels are expressed relative to *ACTIN* as endogenous control using the ΔCt method. Error bars are standard errors of the means based on n = 5. ns = not significant, ** = *P* < 0.01, *** = *P* < 0.001 based on two-tail *t-*tests for each time point. Raw data and statistical analyses are available in Supplemental Data S2.

Although it is not possible to determine heading time in Kronos-PS under SD, the final number of leaves can be used to estimate the timing of the transition from the vegetative shoot apical meristem (SAM) to the reproductive phase (flowering time). In two separate SD experiments, one performed under 8 h of light (SD) and the other under 10 h of light (SD-10h), Kronos-PS primary shoots produced 13.9 and 13.7 leaves in SD and SD-10h, respectively (including leaves still wrapped inside the sheaths). The *gi-2*-PS mutant produced ∼1 more leaf than the Kronos-PS control under both conditions: SD (14.9 leaves) and SD-10h (14.5 leaves, Fig. 2C). However, when the same lines were grown under LD, the difference between wild type and *gi-2* increased to 3.4 leaves, indicating a highly significant interaction (*P<* 0.001) between the effects of *GI* and photoperiod on leaf number (Supplemental Information S2).

This interaction between *GI* and photoperiod was no longer significant when we tested the effect of the *gi-2* mutant on heading time in a Kronos-PI background (henceforth *gi-2-*PI). The *gi-2-*PI mutant showed similar delays in heading time relative to Kronos-PI under both LD (15.4 d) and SD conditions (16.5 d, Fig. 2D). A factorial ANOVA for heading time using these lines, showed highly significant effects for *GI* and photoperiod (*P* < 0.001), but no significant interaction between them (*P* = 0.4347, Supplemental Information S2).

To explore the interactions between *GI* and *PPD1* under LD, we crossed Kronos-PS *gi-2* with a loss-of-function *ppd1* mutant (Pearce et al. 2017) and compared heading times among the four possible homozygous classes. A factorial ANOVA showed highly significant effects of *PPD1* and *GI* on heading times (*P <* 0.001), but no significant interaction between the two genes (Fig. 2E, *P* = 0.3807, Supplemental Information S2). We did not study these lines under SD because *gi-2* and *ppd1* mutants, as well as Kronos-PS controls do not head under SD. Taken together, these results suggest that the *PPD1* PS-allele is important for the *GI* differential flowering responses in LD and SD.

Lastly, we compared *PPD1* expression profiles in Kronos-PS and the *gi-2-*PS mutant under LD (Fig. 2F). In Kronos-PS, *PPD1* transcripts showed diurnal rhythms with a peak expression 12 hours after the lights were turned on (zeitgeber time 12 = ZT12). In the *gi-2-*PS mutant, *PPD1* transcript levels were significantly downregulated relative to Kronos-PS from the ZT12 expression peak to ZT20 (Fig. 2F).

### *GI* and *ELF3* show strong interactions on the regulation of *FT1* expression and heading time

Since loss-of-function mutations in *elf3* greatly reduce the photoperiodic response and significantly alter *PPD1* expression (Alvarez et al. 2016; 2023), we evaluated the effect of *gi-2* in the *elf3* mutant background. In the F_2_ progeny of a cross between *gi-2* and *elf3* (both in Kronos-PS), we selected homozygous *elf3* and *elf3 gi-2* mutants and evaluated them under LD and SD. Both *GI* and photoperiod showed highly significant effects on heading time (*P* < 0.001), however, no interaction was observed between them (*P* = 0.7608, Supplemental Information S3). The delayed heading caused by the *gi-2* mutant in the *elf3* background was similar under LD (14.7 d) and SD (14.9 d, Fig. 3A), indicating that in the absence of a functional *ELF3* the effect of *GI* is no longer modulated by photoperiod.

**Fig. 3.**
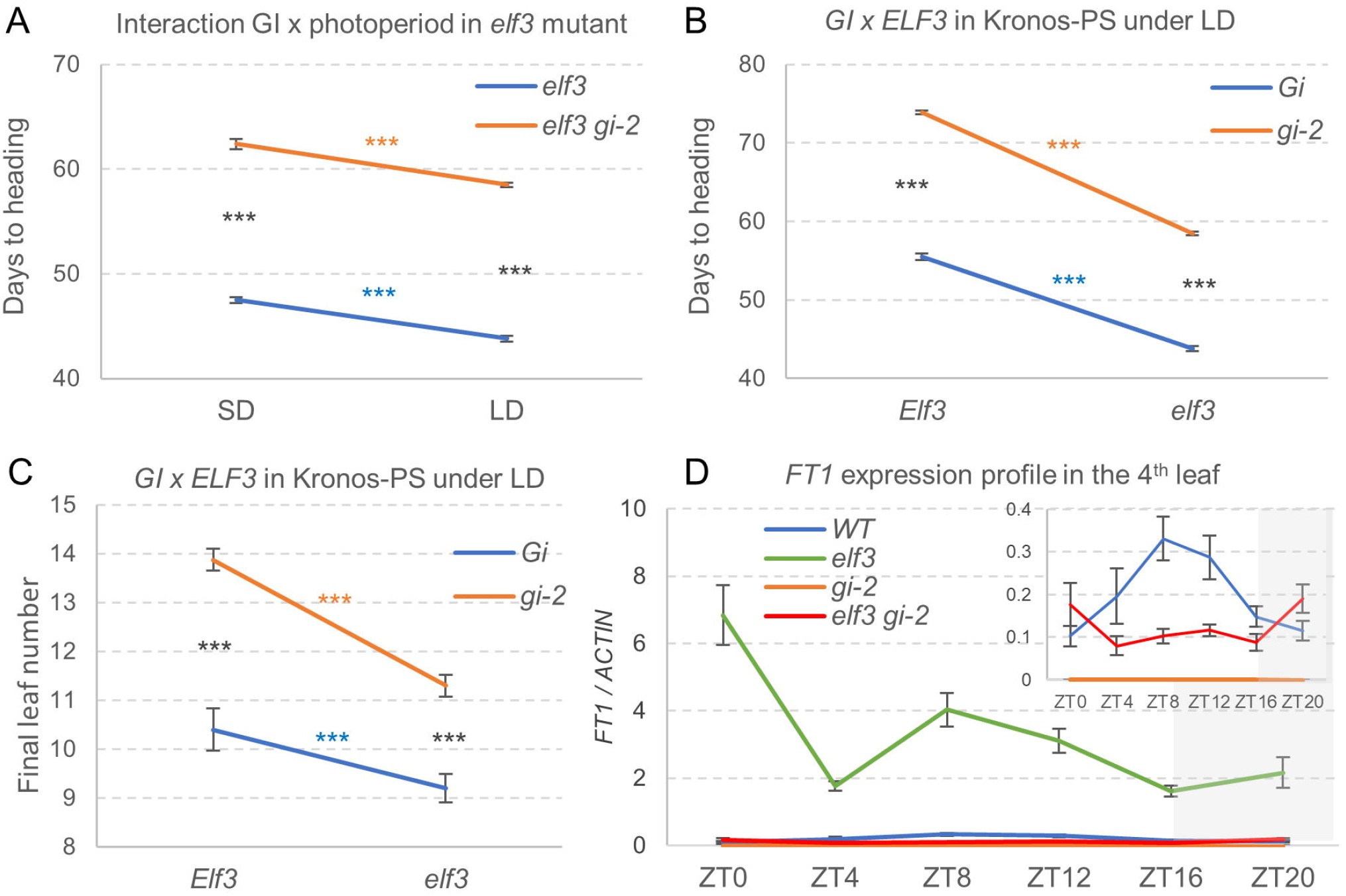
Interactions between *GI* and *ELF3* on the regulation of flowering/heading time and *FT1* expression. A, Interaction graph showing the effect of the *gi-2* mutation on days to heading under different photoperiods in a Kronos-PS-*elf3* mutant background *(elf3* vs. *elf3 gi-2*). SD = 8 h light and LD = 16 h light (n = 10-12). B and C, Interaction graphs for the effects of *GI* and *ELF3* on heading time (B) and total number of leaves produced by the primary shoots (C) under LD. Error bars are standard errors of the means based on n =8-10. *** = *P* < 0.001 for the four simple effect ANOVAs. The 2 x2 factorial ANOVAs are in Supplemental Data S3. D, Time-course expression analysis of *FT1* in the 4^th^ fully expanded leaves of Kronos-PS (WT), *elf3*, *gi-2* and *elf3 gi-2* mutants under LD. The gray shaded area indicates the 8 h of darkness. The inset shows *FT1* expression from the same experiment but excluding the *elf3* mutant to visualize better the smaller differences among the other genotypes. Expression levels were calculated relative to the endogenous control *ACTIN* using the ΔCt method (n = 5). Raw data and statistical analyses are available in Supplemental Data S3.

We further explored the interaction between *GI* and *ELF3* in an additional LD experiment including Kronos-PS, *gi*-2, *elf3*, and *elf3 gi-2* mutants in the same PS genetic background. Consistent with previous findings (Alverez et al. 2016; 2023), the *elf3* allele was associated with a highly significant acceleration in heading time and a reduced number of leaves relative to Kronos-PS, but the differences were larger in the *gi-2* than in the wild-type background (Fig. 3B and C). By contrast, the *gi-2* allele was associated with significant increases in leaf number and delayed heading, with larger differences in the presence of the functional *Elf3* allele than in the *elf3* mutant background (Fig. 3B and C). These differential responses were reflected in highly significant interactions between GI and ELF3 for both traits (*P* < 0.001, Supplemental Information S3).

We next examined the diurnal expression pattern of *FT1* transcripts in the same four genotypes. RNA samples were extracted from the 4^th^ fully expanded leaves (L4, before the SAM transition to the reproductive phase) collected at 4 h intervals for a 24 h period. As expected, the highest transcript levels of *FT1* were observed in the *elf3* mutant (earliest heading) and the lowest in the *gi-2* mutant (latest heading, Fig. 3D). Factorial ANOVAs at each ZT point revealed highly significant effects of *ELF3* and *GI* and highly significant interactions on the regulation of *FT1* (*P* < 0.001, Supplemental Information S3). These results suggest that the differences in heading time and leaf number in the *elf3* and *gi-2* mutants are mediated by their effects and interactions on the regulation of *FT1* expression.

We next explored the effect of the different mutant genotypes on the expression of *GI* and core circadian oscillator genes *CIRCADIAN CLOCK ASSOCIATED1* (*CCA1*, Fig. 4B) and *TIMING OF CAB EXPRESSION 1* (*TOC1*, Fig. 4C). *GI* transcript levels in the *elf3* mutant were significantly lower than in Kronos-PS at the ZT12 peak, but significantly higher at ZT0, ZT4 and ZT20 (Fig. 4A). The diurnal expression patterns of *CCA1* (Fig. 4B) and *TOC1* (Fig. 4C) were altered in the *gi-2* and *elf3* individual mutants, with an even larger disruption observed in the *elf3 gi-2* combined mutant. This was particularly evident in the *CCA1* expression profile, which showed no diurnal rhythms in the combined mutant (Fig. 4B). In summary, these results indicate that the combined *elf3 gi-2* mutant has a very disruptive effect on the circadian clock, a feature that is also conserved in Arabidopsis (Anwer et al. 2020).

**Fig. 4.**
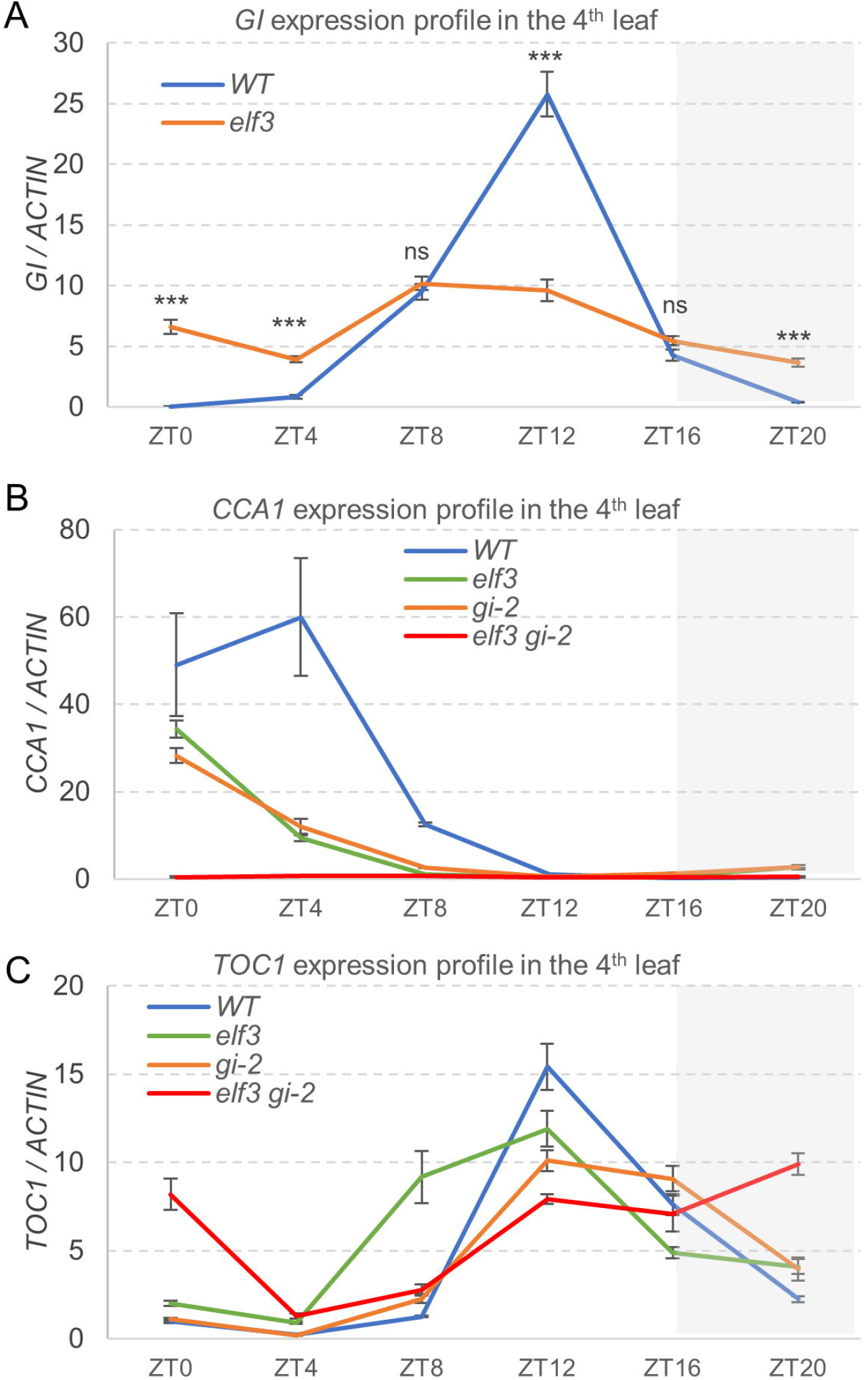
Effect of the *elf3* and *gi-2* mutations on the transcription profile of *GI* and the core circadian oscillator genes. Samples were collected at 4h intervals for a 24h period from the 4^th^ fully expanded leaves. Expression levels of the target genes relative to the *ACTIN* endogenous control were evaluated by qRT-PCR using the ΔCt method. A, Effect of the *elf3* mutation on *GI* transcript levels. *** = *P* < 0.001 based on two-tailed t-test. The shaded gray area indicates the 8 h of darkness. Error bars are s.e.m based on 5 biological replications. B and C, Diurnal expression patterns of the core circadian clock genes *CCA1* (B) and *TOC1* (C) in Kronos-PS (WT), *elf3*, *gi-2,* and combined mutant *elf3 gi-2*. Five biological replicates were used per genotype. Raw data and factorial ANOVAs at each time point are provided in Supplemental Data S4.

Finally, because ELF3 was previously shown to physically interact with GI and modulate the cyclic accumulation of the GI protein in Arabidopsis (Yu et al. 2008), we explored the physical interactions between the wheat GI and ELF3 proteins. Indeed, we observed a strong interaction between the two wheat proteins in yeast two-hybrid (Y2H) assays (Fig. S2). The GI and ELF3 interactions at the protein level, together with the interactions at the transcriptional level, likely contribute to the observed genetic interactions between *GI* and *ELF3* on heading time observed in this study.

### Interactions between *GI* and the LD flowering repressor *VRN2* contribute to the regulation of wheat heading time

To determine the relationship between *GI* and *VRN2*, we crossed *gi-2* with a Kronos *vrn2* loss-of-function mutant (Distelfeld et al. 2009) and studied the effect of the four different homozygous genotypes on heading time. When grown under LD, the *vrn2* mutant headed 5.3 d earlier than the wild type, and this effect increased to 12.4 d when the same comparison was performed in the *gi-2* mutant background (Fig. 5A). On the contrary, the 21.6 d delay in heading time associated with the *gi-2* mutant in plants homozygous for the functional *Vrn2* allele was reduced to 14.5 d in plants homozygous for the *vrn2* mutant allele (Fig. 5A). These differential effects were reflected in a highly significant interaction between *GI* and *VRN2* on heading time (*P* < 0.001, Supplemental Information S5).

**Fig. 5.**
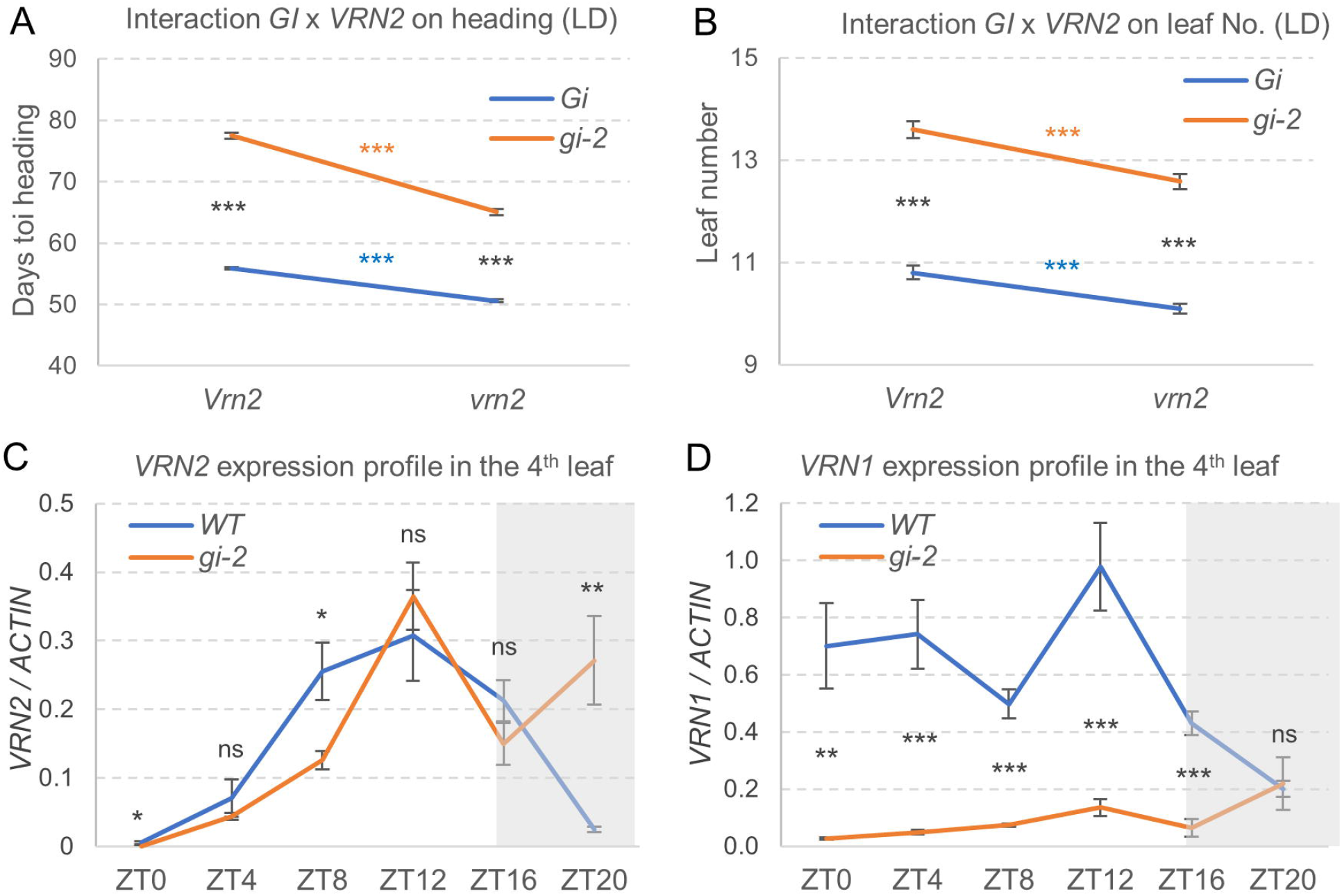
Interactions between *GI* and *VRN2*. A-B, Interaction graphs showing the effect of *GI* on heading time (A) and leaf number (B) in the presence and absence of functional *VRN2* alleles (n = 10-12). C-D, Effect of *gi-2* mutation on the transcript levels of *VRN2* (C) and *VRN1* (D) under LD. Each data point represents the average of 5 biological replicates. Samples were collected at 4h intervals for a 24h period from the 4^th^ fully expanded leaves and transcript levels were evaluated by qRT-PCR and calculated relative to the *ACTIN* endogenous control using the ΔCt method. Error bars are standard errors of the means. ns = not significant, * = *P* < 0.05, ** = *P* < 0.01, and *** = *P* < 0.001 were obtained from simple effect ANOVAS in A and B (complete ANOVA in Supplemental Data S5) and two-tailed t-tests in C and D at each time point. Raw data is available in Supplemental Data S5.

When we analyzed the same plants for total leaf number, we also found highly significant differences associated with *VRN2* and *GI* (Fig. 5B), but the interaction was not significant (*P =* 0.2665, Supplemental Information S5). These results suggest that the interaction between *GI* and *VRN2* has a limited effect on the initial transition of the SAM from the vegetative to the reproductive phase, but a stronger effect on the subsequent phase of spike development and stem elongation.

To characterize better the genetic interaction between *GI* and *VRN2* on heading time, we determined the effect of the *gi* mutation on the *VRN2* transcription profile by qRT-PCR (Fig. 5C). In both the Kronos-PS control and the *gi-2* mutant, *VRN2* transcripts increased during the day to reach a peak at ZT12, with slightly lower levels in *gi-2* at ZT0 and ZT8 (*P<* 0.05). (Fig. 5C). However, after the lights were turned off, *VRN2* transcript levels decreased rapidly in Kronos-PS but remained high in the *gi-2* mutant (Fig. 5C), suggesting a role of *GI* in the downregulation of *VRN2* at night.

Since VRN1 is a known repressor of *VRN2* (Chen & Dubcovsky, 2012), we also analyzed the transcript levels of *VRN1* using the same leaf samples. *VRN1* transcript levels were significantly higher in the wild type than in the *gi-2* mutant in most time points, but at ZT20 the two genotypes showed no significant differences (Fig. 5D). This result suggests that the role of *GI* in the downregulation of *VRN2* during the night is not mediated by *VRN1*, but we cannot completely rule out the possibility of a residual effect of the higher transcript levels of *VRN1* at dusk (ZT16) in Kronos-PS relative to *gi-2*.

In addition to their interactions at the transcriptional level, GI and VRN2 proteins show strong interaction in Y2H assays (Fig. S2). A similar interaction has been reported between the two orthologous rice proteins by Y2H and further validated by BiFC assays in *Nicotiana benthamiana* and by Co-IP experiments in rice protoplasts (Zheng et al. 2019). In summary, the observed interactions between GI and VRN2 at the protein and transcriptional levels likely contribute to their significant genetic interactions on heading time.

### *GI* shows strong interactions with *CO1* and *CO2* on the regulation of heading time in wheat

The regulation of flowering time by *GI* in Arabidopsis is mainly mediated through the *CO*-dependent pathway (Sawa et al. 2007), so we studied the interactions between *GI* and the two *CO* paralogs present in wheat: *CO1* and *CO2* (Shaw et al. 2020). We first determined the effect of the Kronos-PS *gi-2* mutant on the diurnal expression patterns of *CO1* and *CO2* in the 4^th^ leaves. The transcript levels of *CO1* were significantly up-regulated in the *gi-2* mutant relative to the Kronos-PS control at ZT0 and ZT4, but were mostly indistinguishable between the two genotypes at other time points (Fig. 6A). Transcript levels of *CO2* were significantly higher in the *gi-2* mutant relative to Kronos-PS at three (ZT4, ZT8 and ZT12) of the six analyzed time points (Fig. 6B).

**Fig. 6.**
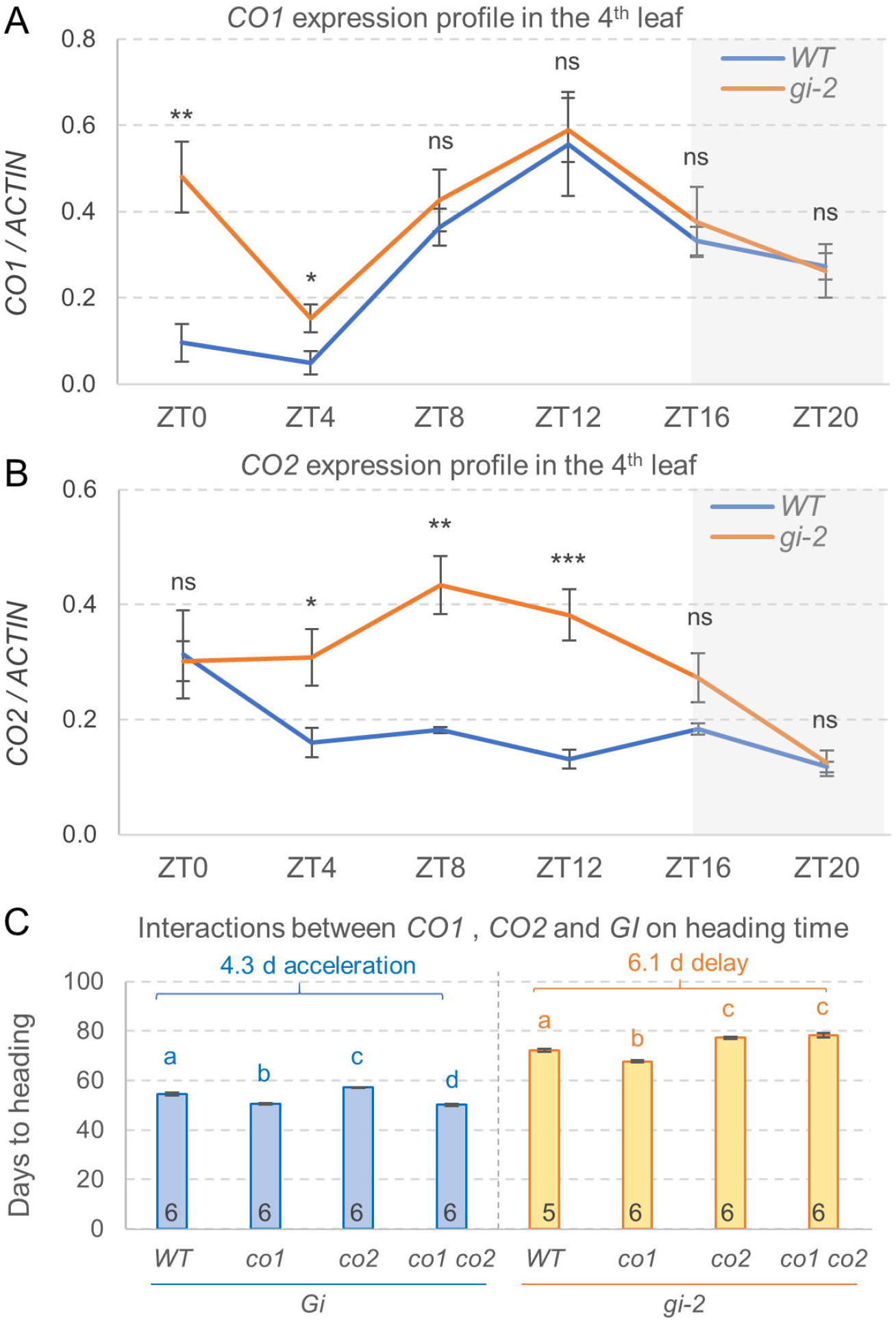
Interactions among *GI, CO1* and *CO2* on the regulation of wheat heading time under LD photoperiod. A-B, Effect of *gi-2* mutation on the expression profiles of *CO1* (A) and *CO2* (B) under LD (16h light). The gray shadowed area indicates the hours of darkness. The 4^th^ fully expanded leaves were collected at 4 h intervals for a 24 h period. Expression levels were calculated relative to the endogenous control *ACTIN* using the ΔCt method. Values are averages of 5 biological replicates and error bars are standard errors of the means. ns = not significant, * = *P* < 0.05, ** = *P* < 0.01, *** = *P* < 0.001 for differences between genotypes at each time point using t-tests. C, Heading time of wild-type Kronos-PS (WT), *co1, co2, co1 co2*, *gi, co1 gi, co2 gi* and *co1 co2 gi* homozygous mutants grown in the same growth chamber under LD. The number of plants included in the experiment are indicated inside the bars. Error bars are standard errors of the means. Different letters with the same color indicate significant differences in Tukey tests (*P* < 0.05) in separate ANOVAs for *Gi* (blue) and *gi-2* (orange) genotypes. Raw data and statistical analysis are available in Supplemental Data S6.

Interestingly, GI also showed strong physical interactions with both CO1 and CO2 proteins in Y2H assays (Fig. S2). These results suggest that wheat GI may also regulate CO protein stability through protein interaction, as previously described in Arabidopsis GI (Song et al. 2014; Hwang et al. 2019). Interactions at both the transcriptional and protein levels may contribute to the genetic interaction between *GI* and *CO1/CO2* on heading time described in the next section.

From the cross between *gi-2* and *co1co2* double mutant in the Kronos-PS background we generated the eight possible homozygous classes and studied their effect on heading time (all genotypes were grown in the same LD growth chamber). Among the four genotypes carrying the *Gi* wild-type allele, *co1* flowered 3.8 d earlier, *co2* 2.7 d later and the *co1co2* combined mutant 4.3 d earlier than Kronos-PS (Fig. 6C), similarly to what was reported in a previous study (Shaw et al. 2020). Among the four genotypes carrying the *gi-2* mutation, the differences between the individual mutants and the wild type (*co1* 4.5 d earlier and *co2* 5.7 d later) were in the same direction as in the corresponding lines with the *Gi* allele (Fig. 6C). However, the *co1 co2* combined mutant was 6.1 d later than the wild type in the *gi-2* background and 4.3 d earlier in the *Gi* background (Fig. 6C). These opposite results were reflected in highly significant interactions among these three genes in a 3-way ANOVA (*P* < 0.001, Supplemental Information S6). The effect of *GI* on heading time was stronger in the presence of the *co1* allele than in the presence of the *Co1* allele, whereas the effect of *CO1* was stronger in *Gi* than in *gi-2* (Fig. 6C). The effects of both *co2* and *gi-2* on heading time were stronger in the presence of the mutant allele of the other gene than in the presence of the wild-type allele (Fig. 6C).

### Interactions between *GI* and the *miR172-AP2* age-dependent pathway contribute to the regulation of *FT1* and wheat heading time

In Arabidopsis, one of the pathways that interacts with *GI* to regulate the expression of *FT* is the age-dependent miR172-*AP2/TOE* pathway (Jung et al. 2007), which is conserved in wheat (Debernardi et al. 2022). To test if a similar interaction exists in wheat, we first determined the effect of *gi-2* (in PS background) on the expression levels of *miR172*, *AP2* and *FT1*. In both L7 and L9 leaves, the mature *miR172* levels were significantly lower in the *gi-2* mutant than in the Kronos-PS control (Fig. 7A). As expected from the known role of *miR172* as a repressor of *AP2LI* (= *TOE1*, Debernardi et al. 2022), the transcript levels of *AP2L1* were higher in the *gi-2* mutant than in the Kronos-PS control (Fig. 7B). Transcript levels of *FT1* were significantly lower in the *gi-2* mutants relative to the wild type at both stages, which is consistent with the known role of *AP2L1* as a repressor of *FT1* (Debernardi et al. 2022) and with the later heading time of the *gi-2* mutant relative to the wild type (Fig. 7C).

**Fig. 7.**
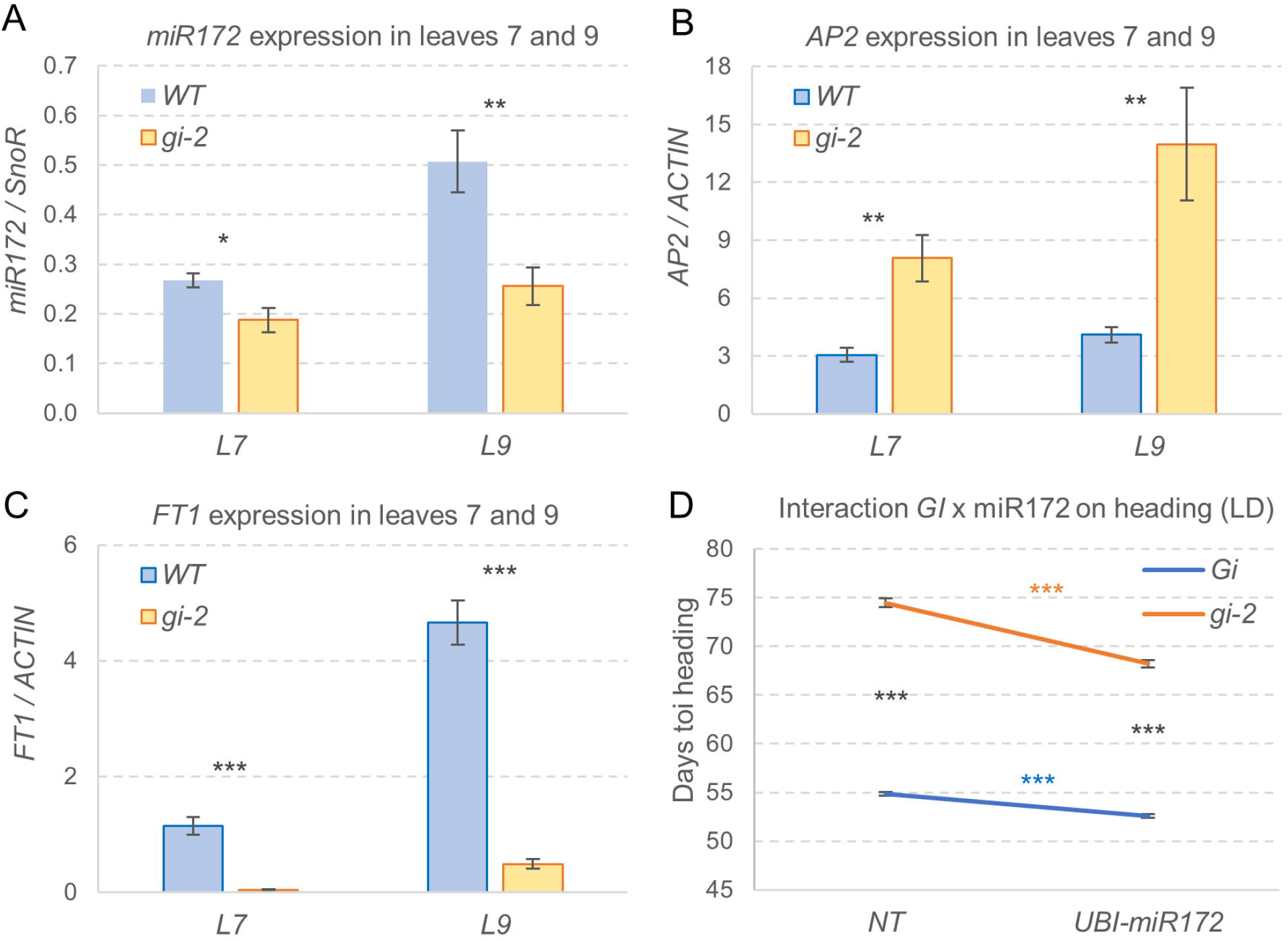
Characterization of interactions between *GI* and *miR172*. A-C, Evaluation of the effect of *gi-2* mutation on the transcript levels of (A) mature *miR172,* (B) *AP2L1*, and (C) *FT1* by qRT-PCR analysis. The mature *miR172* levels were calculated using the small nucleolar RNA 101 (SnoR) as internal reference, and the transcript levels of *AP2L1* and *FT1* were calculated relative to the internal reference *ACTIN* using the ΔCt method. Samples were collected at ZT4 from levels 7 and 9 of Kronos-PS (WT) and *gi-2* mutant plants grown under LD. Values are averages of 5 biological replicates and error bars are standard errors of the means. * = *P* < 0.05, ** = *P* <0.01, *** = *P* < 0.001 indicate differences between the wild type and *gi-2* mutant using two-tail *t-*test. D, Interaction graph showing the effect of the *UBI-miR172* transgene on heading time under LD in the presence and absence of *GI* functional alleles (n = 13-15). *** = *P* < 0.001 indicate the significance of the four simple effect ANOVAs. Raw data and statistical analysis of the complete 2 x2 factorial ANOVA is available in Supplemental Data S7.

To investigate the genetic interaction between miR172 and GI, we crossed the *gi-2* mutant with a *UBI-miRNA172* transgenic line (hereafter *UBI-miR172*) developed previously (Debernardi et al. 2022), and evaluated the four different homozygous genotypes for heading time in a growth chamber under LD. Plants overexpressing *miR172* showed a small but significant acceleration in heading time (Fig. 7D), consistent with a previous study (Debernardi et al. 2022). In the *gi* mutant background, the acceleration of heading time by *UBI-miR172* (6.3 d) was greater than in the wild type (2.3 d), and both were highly significant (Fig. 7D). On the contrary, the delayed heading associated with *gi-2* was smaller in the *UBI-miR172* transgenic plants (15.6 d) than in the non-transgenic sister lines (19.6 d, Fig. 7D). A factorial ANOVA for heading time revealed a highly significant interaction (*P* < 0.001) between *GI* and *UBI-miR172* (Supplemental Information S8). Taken together, these results confirmed that interactions between *GI* and the miR172-*AP2L1* pathway contribute to the regulation of *FT1* expression and to the *GI* effects on wheat heading time.

### Integration of *GI* into a working model of the regulation of wheat heading time

To summarize the multiple pathways through which *GI* regulates wheat heading time, and its interactions with other flowering genes, we present a working model in Fig. 8, and a summary of the protein interactions in Supplemental Fig. S2B. Both figures indicate a highly interconnected regulatory network.

**Fig. 8.**
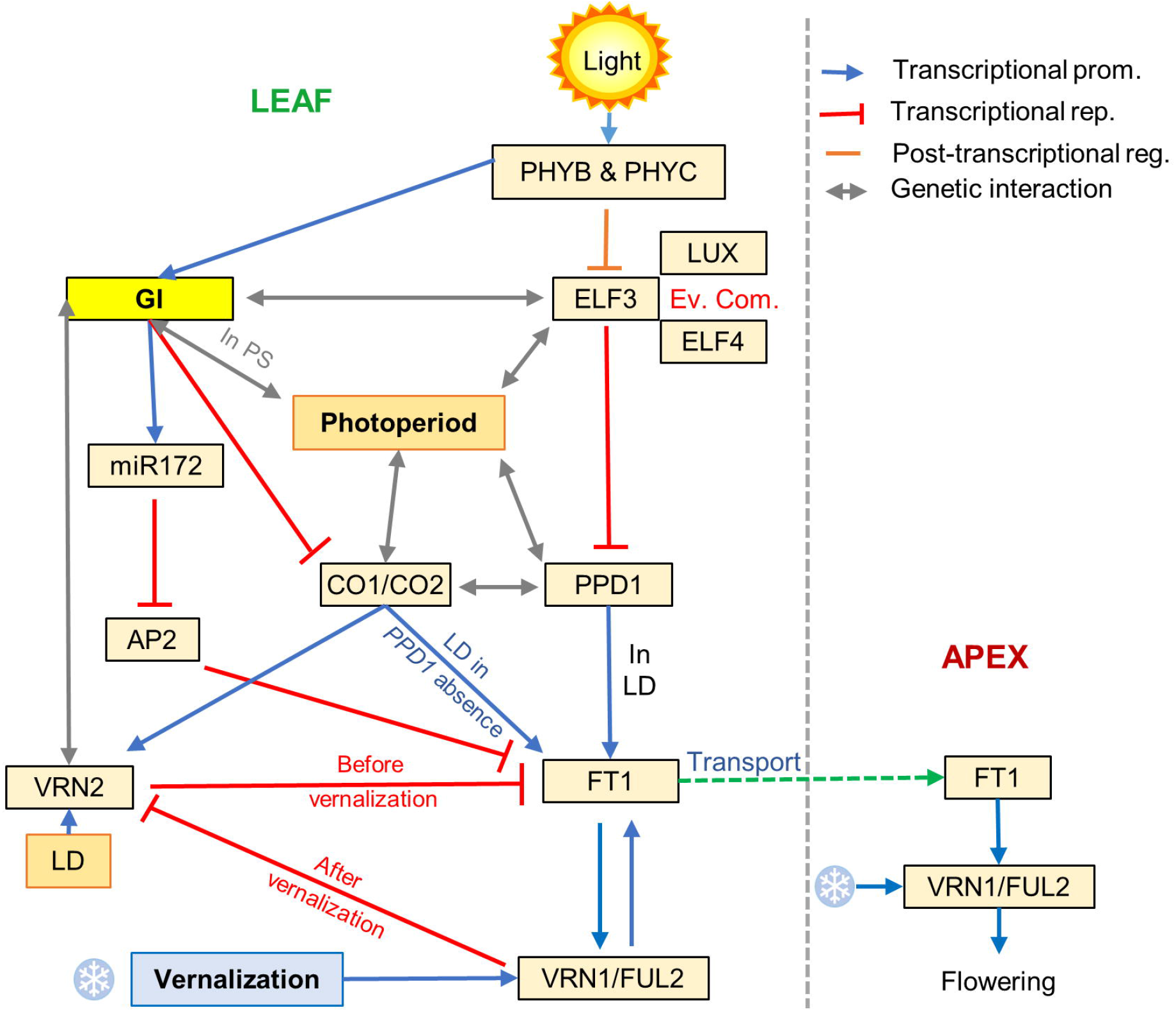
Regulation of heading time in wheat. Blue arrows indicate transcriptional promotion, red lines transcriptional repression, orange lines posttranscriptional regulation, and grey lines genetic interactions.

This model shows that GI promotes *FT1* expression and heading time through multiple pathways, including the grass-specific *VRN2* gene, as well as through complex interactions with *CO1/CO2* and the miR172-AP2 age-dependent pathway. The model also shows that the interactions between *GI* and photoperiod are dependent on the presence of a wild-type *PPD1-*PS allele. Some of the interactions are shown by bidirectional grey arrows because they are too complex to be summarized by a promotion arrow or repression T-bar. For example, *VRN2* expression is upregulated in the *gi-2* mutant at ZT20 but downregulated at ZT0-ZT8 (Fig. 5); and *GI* is downregulated in the *elf3* mutant at ZT12 but upregulated at ZT20-ZT4 (Fig. 4).

This model presents only the strongest effects, but additional weaker effects are known to exist. For example, *PHYB* affects heading time and expression of multiple flowering genes even in the absence of *ELF3*, whereas *ELF3* affects heading time even in the absence of *PPD1* (Alvarez et al. 2023).

## Discussion

### Functional conservation of *GI* homologs in the regulation of flowering time

The role of *GI* as a flowering promoter is conserved among LD plant species grown under inductive LD photoperiods, while some functional diversification can be observed when those plants are grown under non-inductive SD conditions. In Arabidopsis, *GI* activates *FT* expression and promotes flowering time, and most *gi* mutants show delayed flowering phenotypes in both LD and SD conditions, with smaller delays under SD (summarized in Mishra and Panigrahi, 2015). In pea, mutations in *LATE BLOOMER1* (*LATE1*, *GI* homolog in pea) prevent the induction of *FT* and result in late flowering under LD (Hecht et al. 2007), confirming the function of *LATE1* as a LD flowering promoter. In photoperiod sensitive wheat genotypes, the promoting effect of *GI* on heading time is also stronger under LD than under SD (Fig. 2C).

*GI* homologs from SD grass species, such as rice and sorghum, promote flowering under SD photoperiods, with *gi* mutants showing reduced florigen expression and late flowering under this condition. However, under LD photoperiods rice *gi* mutant shows no changes in heading time (Izawa et al. 2011), whereas sorghum *gi* mutant still shows delayed flowering (Abdul-Awal et al. 2020). Although the *GI* promotion of heading in SD grasses seems opposite to what we described above for LD species, in both groups, *GI*’s stronger promotion of flowering occurs under the inductive photoperiods (LD for LD-plants and SD for SD-plants). This result is not surprising, since *GI* promotes flowering though multiple pathways that converge on the upregulation of *FT*, which is expressed under inductive photoperiods. *GI’*s stronger photoperiodic response under inductive photoperiods is conserved in LD species from wheat (Fig. 8) to Arabidopsis (Araki and Komeda, 1993; Fowler et al. 1999), and SD species from rice (Izawa et al. 2011) to sorghum (Abdul-Awal et al. 2020), suggesting a conserved role of *GI* in the photoperiodic response.

### The interaction between *GI* and photoperiod is disrupted in Kronos PI and the *elf3* mutant

Grass species have a unique *PPD1*/*PRR37*-dependent photoperiodic flowering pathway in addition to the *CO*-dependent pathway, but their relative importance varies across species (Yano et al. 2000; Koo et al. 2013; Murphy et al. 2011; Yang et al. 2014; Pearce et al. 2017; Shaw et al. 2020). In wheat and barley, *PPD1* plays a more dominant role than *CO1* and *CO2* in the photoperiodic regulation of heading time (Alqudah et al. 2014; Shaw et al. 2020). Most of the natural variation in the photoperiodic response in wheat is associated with either deletions in the promoters of *PPD-A1* or *PPD-D1* or with copy number variation of *PPD-B1* (Wilhelm et al. 2009; Beales et al. 2007; Díaz et al. 2012; WuLJrschum et al. 2019). The deletion in the promoter of *PPD1* results in misexpression of *PPD1* during the night and dawn, and the induction of *FT1* and earlier heading under SD (Alvarez et al. 2023; Wilhelm et al. 2009).

The misexpression of *PPD1* in Kronos-PI also resulted in the elimination of the significant interaction on leaf number observed in Kronos-PS between *GI* and photoperiod (Fig. 2C). These results indicate that the photoperiod sensitive *PPD1* allele is critical for conveying the photoperiodic information to *GI* and for generating its differential flowering response in LD and SD (Fig. 2C).

This hypothesis is further supported by the absence of a significant interaction between *GI* and photoperiod on heading time in the *elf3* mutant, where the expression profiles of both *GI* (Fig. 4A) and *PPD1* (Alvarez et al. 2023) are significantly disrupted. In the absence of *ELF3* function, *GI* can still promote flowering time, although its effect is weaker in *elf3* relative to the *ELF3* wild-type allele (Fig. 3B-C). Moreover, in the absence of *ELF3*, the effect of *GI* on flowering is similar in LD and SD, as *elf3 gi* mutant show similar delays in heading time relative to the *elf3* mutant under both photoperiods (Fig. 3A). These results indicate that the differential effect of *GI* on heading time in different photoperiods is also dependent on a functional *ELF3*.

The altered *PPD1* expression in *elf3* may contribute to the lost interaction between *GI* and photoperiod, but additional post-translational mechanisms cannot be ruled out. In this study, we show that GI and ELF3 physically interact with each other by Y2H assay (Fig. S2), an interaction that has been also observed in Arabidopsis (Yu et al. 2008). In Arabidopsis, GI protein stability and photoperiodic flowering is also regulated by the interaction between ELF3 and COP1 (Yu et al. 2008). ELF3 is essential for the interaction between GI and COP1, and the Arabidopsis *elf3* mutant exhibits a disruption in the cyclic accumulation of GI associated with early flowering and photoperiod-insensitivity (Zagotta et al. 1996; Yu et al. 2008). Similar protein interactions may also contribute to the disrupted photoperiodic response of *GI* in the *elf3* wheat mutant background.

### *GI* interacts with the grass-specific *VRN2* repressor to regulate wheat heading time

The effects of *VRN2* on heading time reported in this study are smaller than those reported in previous studies in winter wheat (Yan et al. 2004; Distelfeld et al. 2009). In winter cereals, *VRN2* is a LD flowering repressor that prevents *FT1* expression during the fall, and is repressed in the spring by *VRN1*, which is upregulated by vernalization (Chen & Dubcovsky, 2012). However, in spring wheats, such as the Kronos variety used in this study, *VRN1* induction does not require vernalization and the earlier down-regulation of *VRN2* results in smaller effects of *VRN2* on heading time in spring than in winter wheat varieties.

In this study, we show that the effects of *VRN2* on heading time are modulated by its interactions with *GI* (Fig. 5A-B) both at the transcriptional and protein levels. In the *gi-2* mutant, the expression levels of *VRN2* vary during the day, with significant increases at ZT20 and significant decreases between ZT0 and ZT8 (Fig. 5C). These diurnal changes likely reflect *GI*’s own circadian oscillation (Fig. 4A) and/or the effects of the *gi-2* mutants on the central oscillators (Fig. 4B-C). GI and VRN2 also interact at the protein level (Fig. S2), a result that has been validated for the rice orthologs (Zheng et al. 2019). A recent study in rice has shown that co-expression of GI and GHD7 causes reduced accumulation of the GHD7 protein, and that GI and phytochromes function antagonistically in the regulation of GHD7 protein stability (Zheng et al. 2019).

The existence of a similar negative effect of GI on the stability of the VRN2 protein in wheat is indirectly supported by the stronger effect of *VRN2* on heading time in the *gi-2* mutant than in the presence of the functional *Gi* allele (Fig. 5A-B). This significant genetic interaction between *GI* and *VRN2* on heading time also resulted in larger differences between the *gi-2* mutant and the wild type in the presence of the functional *Vrn2* allele than in the *vrn2* mutant. This result indicates that part of the effect of *GI* in the acceleration of heading time in wheat is mediated by its interactions with *VRN2*.

In summary, these results show that in the grasses, GI interacts with *VRN2*, a gene that has no homologs in the eudicots plants. It will be interesting to determine if these genetic interactions are the result of direct interactions between these two genes or an indirect effect of the multiple interactions between *GI* and other flowering genes.

### The conserved GI-CO pathway contributes to the regulation of wheat heading time

The mechanistic characterization of *GI*’s role in photoperiodic flowering regulation is mostly from Arabidopsis, where GI promotes *CO* transcription and acts as a major mediator between the circadian clock and CO. Under LD, the peak of GI protein accumulation coincides with FKF1 in the middle of the day, resulting in the repression of CDF1 and, consequently, in increased transcript levels of *CO* and *FT* (Sawa et al. 2007). Under SD, GI accumulation peaks three hours earlier than that of FKF1, precluding the repression of CDF1, and resulting in reduced levels of *CO* transcripts (Sawa et al. 2007). Therefore, the formation of the GI-FKF1 complex is required for day-length measurement in Arabidopsis. In addition, GI can regulate CO temporal stability through direct physical interactions and through its interactions with FKF1 and ZTL (Song et al. 2014; Hwang et al. 2019).

In this study, we show a direct physical interaction between GI and both CO1 and CO2 wheat proteins in Y2H assays (Fig. S2). A similar interaction between GI and CO has been reported in Arabidopsis (Song et al. 2014), suggesting similar mechanisms between the two species. This hypothesis is also supported by the physical interactions observed between GI and ZTL, COP1 and FKF1 in *Brachypodium* (Hong et al. 2010) and GI-FKF1 interaction in sorghum (Abdul-Awal et al. 2020). The physical interactions between GI and both CO1 and CO2 in wheat are reflected in a strong genetic interaction, where the differences in heading time between Kronos-PS and the combined *co1 co2* mutant are reverted in the *gi-2* mutant relative to the wild type (Fig.6C).

We detected differences between wheat and Arabidopsis in the transcriptional regulation of *CO* by GI. While transcript levels of *CO* are significantly reduced in the Arabidopsis *gi* mutant relative to the wild type (Sawa et al. 2007; Sawa and Kay, 2011), *CO1* and *CO2* are significantly upregulated at different times of the day in the wheat *gi-2* mutant (Fig. 6A-B). Therefore, although the presence of the GI-CO pathway seems to be conserved in Arabidopsis and wheat, the mechanisms by which they control photoperiodic flowering likely differ between these two species.

These differences are also reflected in the more limited role of *CO1* and *CO2* in the regulation of the photoperiodic response in the temperate grasses, where *PPD1* is the main photoperiod gene (Shaw et al. 2020). Although both the *PPD1*-dependent pathway and the *CO*-dependent pathway can perceive the differences in photoperiod in the absence of the other gene, there are still significant genetic interactions among them that are important for the fine-tuning of the flowering response in wheat (Shaw et al. 2020).

### GI can regulate wheat heading time by CO-independent regulatory pathways

In addition to the CO-mediated pathway, we show that in wheat GI can also regulate *FT1* expression through the age-dependent miR172-AP2 module. The interaction between GI and miR172-AP2 seems to be conserved between Arabidopsis and wheat, with GI promoting miR172 and repressing AP2 expression in both species (Jung et al. 2007; Fig. 7).

In Arabidopsis, GI has been shown to promote *FT* transcription through direct binding to its promoter regions (Sawa & Kay, 2011). However, as SVP, TEM1 and TEM2 also bind to the same *FT* promoter regions, it is also possible that the binding of GI to the *FT* promoter is mediated by a complex with other proteins (Sawa & Kay, 2011). In wheat, we also observed a strong downregulation of *FT1* in the *gi-2* mutant (Fig. 7C), which was observed even in the presence of the *elf3* mutation (Fig. 3D). Although these results may reflect a direct interaction between GI and the *FT1* promoter, we cannot rule out indirect effects mediated by the multiple pathways through which GI regulates *FT1*.

## Materials and methods

### Plant materials

The reference sequences of *GI* (TraesCS3A02G116300 and TraesCS3B02G135400) were used to screen for mutations in the sequenced ethyl methanesulfonate mutagenized Kronos population (Krasileva et al. 2017) using BLASTN. A splicing site mutation (*A-401*) and three stop codon mutations (*A-2019*, *B-2205* and *B-3825*) were selected to generate *GI* loss-of-function mutants. To test genetic interactions between *GI* and *ELF3*, *CO1*, *CO2*, *PPD1* and *VRN2*, we combined *gi-2* with the respective loss-of-function mutants of these genes and generated *elf3 gi*, *co1 gi*, *co2 gi*, *co1 co2 gi*, *ppd1 gi* and *vrn2 gi* in the same Kronos-PS background. Primers used to genotype and verify the presence of individual gene mutations are listed in Table S1. We also crossed the *gi-2* mutant with a *Ubi-miR172* transgenic line (Debernardi et al. 2022) to investigate the interaction between miR172 and *GI* in the Kronos-PS background. Primers used to differentiate the *Ppd-A1b* allele (PS) from *Ppd-A1a* allele (PI) are also included in Table S1.

### Growth conditions

All experiments were performed in controlled environment conditions using the Conviron PGR15 growth chambers. During the lights-on period, the growth chambers were set at 22°C, but the first and last hour of this lights-on period were set at 20°C to provide a more gradual change between temperatures. Night temperatures were set at 17°C. All PGR15 chambers used similar metal halide and high-pressure sodium light configurations, and lights were set to the same intensity in all experiments (∼330µM m^-2^ s^-1^). Lights were on for 8 hours for SD-8h, 10 hours for SD-10h experiments and 16 hours for LD experiments.

### Generation of the transgenic line overexpressing *GI*

Transgenic Kronos plants overexpressing *GI* were generated at the UC Davis Plant Transformation Facility (http://ucdptf.ucdavis.edu/) using the Japan Tobacco vector pLC41 (hygromycin resistance) and transformation technology licensed to UC Davis. The coding region of *GI-A* gene from Kronos was cloned downstream of the maize UBIQUITIN promoter with a C-terminal 4xMYC tag. Agrobacterium strain EHA105 was used to infect Kronos immature embryos and all transgenic plants were genotyped by PCR using primers described in Supplemental Table S2.

### qRT-PCR analysis

For the expression analyses across developmental stages, we collected the newest fully expanded leaf samples (L7, L9 and L11) 4 hours after lights were turned on (at ZT4). For the time-course experiments analyzing the diurnal gene expression patterns, the 4^th^ fully expanded leaves were collected at 4h intervals for 24 hours. RNA samples were extracted using the Spectrum Plant Total RNA Kit (Sigma-Aldrich). Total RNA was treated with RQ1 RNase-free DNase (Promega) prior to cDNA synthesis. And 1µg of total RNA was used to synthesize cDNA using the High-Capacity cDNA Reverse Transcription Kit (Applied Biosystems) according to the manufacturer’s instructions. The cDNA was then diluted 20 times and 5 µl of the dilution was mixed with 2×VeriQuest Fast SYBR Green qPCR Master Mix (Affymetrix, 75690) and primers for the real-time qPCR analysis. Primer sequences are listed in Table S2. *ACTIN* was used as endogenous control except for miR172, the small nucleolar RNA 101 (SnoR) was used as internal reference. Primers used in qRT-PCR analyses are included in Table S2.

## Acknowledgments and Funding

This project was supported by the Howard Hughes Medical Institute (https://www.hhmi.org/) and by competitive Grants 2016-67013-24617 and 2022-68013-36439 from the United States Department of Agriculture, National Institute of Food and Agriculture (https://nifa.usda.gov/).

## Author contributions

CL and HL designed and performed most of the experiments and analyses. CL wrote the first version of the manuscript. JMD contributed the *UBI-miR172* transgenic material and determined the effect of GI on the transcript levels of *miR172* and *AP2*. CZ contributed to the diurnal gene expression analyses. All authors reviewed the manuscript, contributed ideas and provided valuable suggestions. CL and JD were responsible for the conceptualization of the project. JD was responsible for obtaining funding, project coordination and supervision, statistical analyses and for the final revision of the manuscript.

## Conflict of interest

The authors declare no conflict of interest

## Data and materials availability

All Kronos mutants and transgenic lines are available from the authors upon requests without any restrictions for use. Kronos single mutants are also available from the Germplasm Resources Unit (GRU) at the John Innes Centre.

## Supplemental Materials

**Fig. S1**. Expression analysis of the transgenic *UBI-GI-MYC* lines (in Kronos-PI background) and evaluation of the effect of the transgene on heading time under long-(LD) and short-day (SD).

**Fig. S2**. GI protein interacts with ELF3, CO1, CO2 and VRN2, but not with PPD1 in Y2H assays.

**Table S1**. Primers used for mutant genotyping.

**Table S2**. Primers used in the qRT-PCR analysis.

**Supplemental Information** S1-S8: raw data and statistics supporting figures.

## Notes

### Competing Interest Statement

The authors have declared no competing interest.

### Summary of Updates

We incorporated supplemental Tables S1 and S2, Figures S1 and S2, and supplemental information S1-S8.

