## Supplemental Figures and Tables for "*GIGANTEA* accelerates wheat heading time through gene interactions converging on *FLOWERING LOCUS T1*"

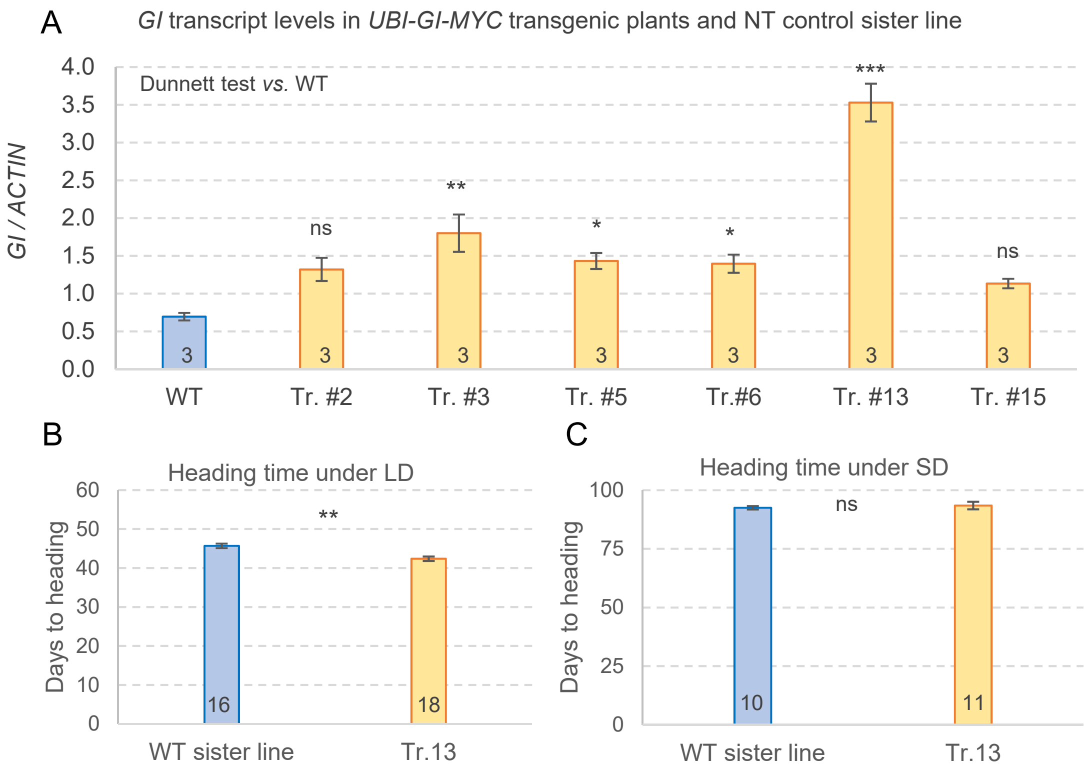


**Fig. S1**. Expression analysis of the transgenic *UBI-GI-MYC* lines (in Kronos-PI background) and evaluation of the effect of the transgene on heading time under long- (LD) and short-day (SD). A, qRT-PCR analysis of the *GI* transcript levels relative to *ACTIN* in six independent transgenic lines constitutively expressing *GI-MYC* under the control of maize Ubiquitin promoter. Tr = Transgenic, WT = Non-transgenic sister-line, n = 3 biological replications. *P* values are two-tailed Dunnett tests of each independent event against the non-transgenic sister line. B, Transgenic line Tr. #13 (Kronos-PI background) headed 3.3 days later than the non-transgenic sister lines under LD. C, Same transgenic line as in (B) showed no significant differences with the control under SD. Numbers inside the bars indicate the number of biological replicates. Error bars are standard errors of the means. For B and C, *P* values are two tail *t*-Tests. ns = not significant, * = $P < 0.05$ ** = $P < 0.01$, and *** = $P < 0.001$. Raw data and statistical analyses are available in Supplemental Data S8.

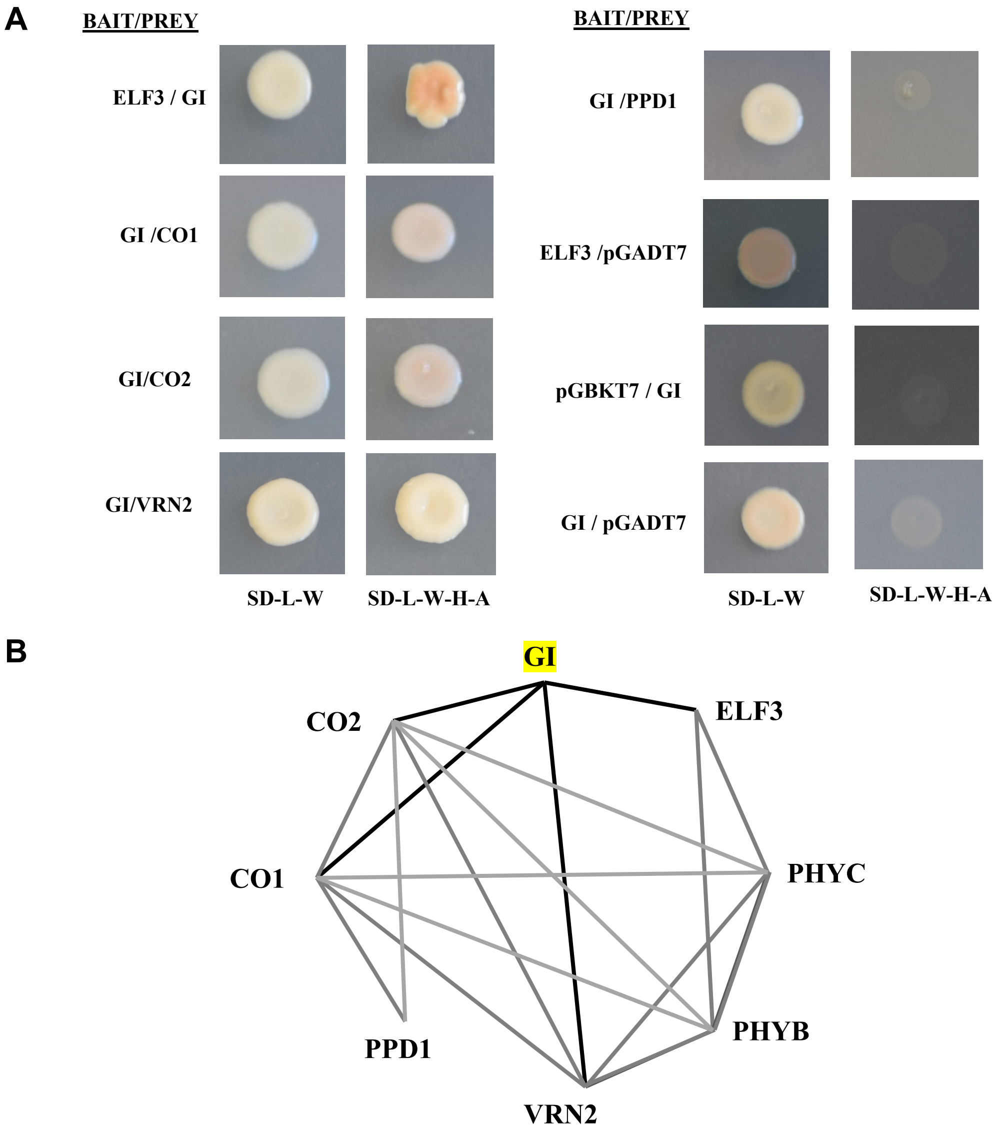


**Fig. S2**. GI protein interacts with ELF3, CO1, CO2 and VRN2, but not with PPD1 in Y2H assays. A, For interaction between ELF3 and GI, the ELF3 full-length protein was fused to GAL4 DNA-binding domain and used as bait and the full-length GI was fused to GAL4 activation domain and used as prey. For interactions with the other proteins, GI protein was used as bait, and CO1, CO2, VRN2 and PPD1 were used preys. SD medium lacking Leucine and Tryptophan (-L-W) was used to select for yeast transformants containing both bait and prey vectors. Interactions were determined on SD media lacking Leucine, Tryptophan, Histidine and Adenine (-L-W-H-A). B, Summary of Y2H interactions among GI reported in this study (black lines) and previously reported interactions (grey lines).

**Table S1**. Primers used for mutant genotyping

| Mutant | Primer sequences | | Marker type and restriction enzyme |
|---|---|---|---|
| ***gi-A-401*** | Common Forward | GTTCTGAGGTGAGGCCCTTACTTTT | KASP |
| | Reverse-1 | GAAGGTGACCAAGTTCATGCTGGCCCACTGCTCTGAGTACTCTTTC | |
| | Reverse-2 | GAAGGTCGGAGTCAACGGATTGGCCCACTGCTCTGAGTACTCTTTT | |
| ***gi-A-2019*** | Common Forward | TGAGATTTCTACCAGGGCATCATCC | KASP |
| | Reverse-1 | GAAGGTGACCAAGTTCATGCTTGGTGCCATTTTCCTCAATGTGTTG | |
| | Reverse-2 | GAAGGTCGGAGTCAACGGATTTGGTGCCATTTTCCTCAATGTGTTA | |
| ***gi-B-2205*** | Forward-1 | GAAGGTGACCAAGTTCATGCTCTTCACTGAAGCGATGTAAGTGG | KASP |
| | Forward-2 | GAAGGTCGGAGTCAACGGATTCTTCACTGAAGCGATGTAAGTGA | |
| | Common Reverse | CATGCAAGTCGATCAAATGGTAGAGA | |
| ***gi-B-3825*** | Common Forward | GACACCTGATGTTAATGCTAT | KASP |
| | Reverse-1 | GAAGGTGACCAAGTTCATGCTTTGCTCGCTATCAGATGTGAACTG | |
| | Reverse-2 | GAAGGTCGGAGTCAACGGATTTTGCTCGCTATCAGATGTGAACTA | |
| ***vrn2*** | Forward | AACGCTTTATGATGCCAAGG | CAPS for linked SNF2 HpyCh4IV[1] |
| | Reverse | TGTGGACAGAACTGGTTTGC | |
| ***ppd-A1*** | Forward | AACGAGCTTAAGAACCACTG | dCAPS, BsrI[2] |
| | Reverse | TATAATAATCACACACGTTG | |
| ***ppd-B1*** | Forward | gcgtaagttactatctctcatggtgtatc | Taqman[3] |
| | Reverse | tttgttttagtacccagtaccataccag | |
| | Probe | [FAM]-ctgctgcttcagttcctagtttcacttgtgtcc-[BHQ1] | |
| ***Ppd-A1a*** | Forward-1 | GTATGCGATTCGCCTGAAGT | Differentiate *Ppd-Ala* and *Ppd-A1b* alleles[4] |
| | Forward-2 | CGTCACCCATGCACTCTGTT | |
| | Reverse | CTGGCTCCAAGAGGAAACAC | |
| ***co-A1*** | Forward | ACATAGGCAGTGCATGAACACAT | CAPS, Hpy188I[5] |
| | Reverse | AGAAGTAGAAAAAGTTGAAGAAAGAG | |
| ***co-B1*** | Forward | CAATTCATCTCTAGGAAAGTAA | CAPS, EcoRV[5] |
| | Reverse | CGTGCTATCTGAAACTATAAAT | |
| ***co-A2*** | Forward | GGTTACAACTCTGGATGGTAG | CAPS, EcoRV[5] |
| | Reverse | GGACTATGTGGTTCACAATATG | |
| ***co-B2*** | Common Forward | GAGGAAGTGGACTCTTGGCTCCTT | KASP (this study) |
| | Reverse-1 | GAAGGTGACCAAGTTCATGCTCGATGTACCTGTTCTGCACATGCTG | |
| | Reverse-2 | GAAGGTCGGAGTCAACGGATTCGATGTACCTGTTCTGCACATGCTA | |
| ***elf3-A*** | Common Forward | GCTTCACCATCTCAAGACAATGAT | KASP[6] |
| | Reverse-1 | GAAGGTGACCAAGTTCATGCTTGAGGCGGAGGAGCACAC | |
| | Reverse-2 | GAAGGTCGGAGTCAACGGATTGAGGCGGAGGAGCACAT | |
| ***elf3-B*** | Common Forward | CTTCACCATCTCAAGACAATGAC | KASP[6] |
| | Reverse-1 | GAAGGTGACCAAGTTCATGCTGGAGCACACCAGTTGTTCTG | |
| | Reverse-2 | GAAGGTCGGAGTCAACGGATGGAGCACACCAGTTGTTCTA | |

**Table S2**. Primers used in the qRT-PCR analysis

| Gene | Primer sequences | |
|---|---|---|
| ***GI*** | Forward | ACTGCACCTTTGGGCATTAG |
| | Reverse | GGCTGTAAGCAGTTGTTGGAG |
| ***FT1***[1] | Forward | CAGCAGCCCAGGGTTGAG |
| | Reverse | ATCTGGGTCTACCATCACGAGTG |
| ***CCA1***[2] | Forward | CCTGGAATTGGAGATGGAGA |
| | Reverse | TGAGCATGGCTTCTGATTTG |
| ***TOC1***[3] | Forward | GCTCATACGCCACCAAGA |
| | Reverse | ACCACACATTCCGCGT |
| ***CO1***[4] | Forward | CACATCAGAGTGGTTATGC |
| | Reverse | GGACTGGACCGTATTGTC |
| ***CO2***[4] | Forward | AAGGGTGTGAGTGTGTAG |
| | Reverse | GATATGTCATTGCTGATGGAAG |
| ***PPD1***[4] | Forward | CGGCATTCACGAGGTACAATAC |
| | Reverse | GAGCCTTGCTTCATCTGAGCG |
| ***VRN2***[5] *(ZCCT2)* | Forward | CCACCATCGTGCCATTCT |
| | Reverse | CCCACCATCATCTCTGTATCAA |
| ***VRN1***[1] | Forward | AAGAAGGAGAGGTCACTGCAGG |
| | Reverse | GGCTGCACTGCCGCA |
| ***Actin***[5] | Forward | ACCTTCAGTTGCCCAGCAAT |
| | Reverse | CAGAGTCGAGCACAATACCAGTTG |
| ***AP2L1***[6] | Forward | ACAACCCATGCCACTCTTCT |
| | Reverse | GTGTTGGTTGCTTGACGATG |
| ***miR172***[6] | SLO-miR172 (RT) | GTCTCCTCTGGTGCAGGGTCCGAGGTATTCGCACCAGAGGAGACATGCAG |
| | Uni-MTRs | TGGTGCAGGGTCCGAGGTATT |
| | RT-miR172 | GGCGGAGAATCTTGATGATG |
| ***snoR101***[6] | Forward | GATGTCTTACACTTGATCTCTGAACTT |
| | Reverse | TGCATCAGGATTGATATAGTGTCC |
